# Brain-age in midlife is associated with accelerated biological aging and cognitive decline in a longitudinal birth-cohort

**DOI:** 10.1101/712851

**Authors:** Maxwell L. Elliott, Daniel W. Belsky, Annchen R. Knodt, David Ireland, Tracy R. Melzer, Richie Poulton, Sandhya Ramrakha, Avshalom Caspi, Terrie E. Moffitt, Ahmad R. Hariri

**Author notes:** Correspondence: Ahmad R. Hariri, Ph.D., Professor of Psychology and Neuroscience, Director, Laboratory of NeuroGenetics, Head, Cognition and Cognitive Neuroscience Training Program, Duke University, Durham, NC 27708, USA.

## Abstract

An individual’s brain-age is the difference between chronological age and age predicted from machine-learning models of brain-imaging data. Brain-age has been proposed as a biomarker of age-related deterioration of the brain. Having an older brain-age has been linked to Alzheimer’s, dementia and mortality. However, these findings are largely based on cross-sectional associations which can confuse age differences with cohort differences. To illuminate the validity of brain-age a biomarker of accelerated brain aging, a study is needed of a large cohort all born the same year who nevertheless vary on brain-age. In a population-representative 1972-73 birth cohort we measured brain-age at age 45, as well as the pace of biological aging and cognitive decline in longitudinal data from childhood to midlife (N=869). In this cohort, all chronological age 45 years, brain-age was measured reliably (ICC=.81) and ranged from 24 to 72 years. Those with older midlife brain-ages tended to have poorer cognitive function in both adulthood and childhood, as well as impaired brain health at age 3. Furthermore, those with older brain-ages had an accelerated pace of biological aging, older facial appearance and early signs of cognitive decline from childhood to midlife. These findings help to validate brain-age as a potential surrogate biomarker for midlife intervention studies that seek to measure treatment response to dementia-prevention efforts in midlife. However, the findings also caution against the assumption that brain-age scores represent only age-related deterioration of the brain as they may also index central nervous system variation present since childhood.

## Introduction

While old age is associated with higher risk for disease across the entire body, degeneration of the brain and consequent cognitive decline has an outsized influence on disability and loss of independence in older adults^1^. As such there is growing need for interventions to slow the progression of cognitive decline. Unfortunately, to date, tested interventions have not slowed age-related cognitive decline^2^. The failure of these interventions may be related to their targeting of individuals too late in the aging process after neurodegeneration has become inexorable^3,4^. Alzheimer’s disease and related dementias (ADRD) arise at the end of a chronic pathophysiological process with preclinical stages emerging decades earlier in life^3^. Evaluating interventions to prevent ADRD onset requires the identification of surrogate biomarkers that index subclinical cognitive decline, neurodegeneration, and accelerated aging of the brain by midlife.

While everyone ages chronologically at the same rate, this is not true biologically; some individuals experience accelerated age-related biological degeneration^5,6^. For decades, researchers have worked to quantify the rate of biological aging and better understand the mechanisms that generate individual differences in the aging process^7^. The resulting measures of accelerated biological aging have been associated with health span, cognitive decline, cancer risk, and all-cause mortality^5,6,8^. However, such aging biomarkers have not directly quantified aging in the organ most directly linked to ADRD, namely the brain. To address this gap, a recently developed measure called “brain-age” has been proposed as a biomarker for accelerated aging of the brain^9,10^. Brain-age is a relatively novel measure derived from neuroimaging, but its interpretation is uncertain.

Brain-age is estimated by training machine-learning algorithms to predict age from structural magnetic resonance imaging (MRI) data collected in large samples of individuals across a broad age range^11^. These machine-learning algorithms “learn” multivariate patterns from MRI data that are useful in explaining variance in chronological age across individuals. These algorithms can then be applied to independent samples to predict chronological age in individuals based on their brain-imaging data. The difference between an individual’s predicted age based on MRI data and their chronological age (hereafter “brain-age”) is usually interpreted as a measure of accelerated aging of the brain. Older brain-age has been associated with mild cognitive impairment, ADRD, and mortality^11,12^. Individuals with an older brain-age are more likely to have risk factors for dementia including obesity, diabetes, alcoholism, and traumatic brain injury^9,12–14^. Initial studies suggest that brain-age may be able to predict cognitive decline and conversion to ADRD in older adults in their 60s, 70s and 80s^15,16^. But there is no evidence linking brain-age to earlier signs of cognitive decline or accelerated aging in midlife, the age when surrogate biomarkers may be more effectively used in ADRD-prevention efforts^4^. Promising results notwithstanding, research on brain-age is still in its infancy. Reported associations between brain-age and risk factors for accelerated aging are largely cross-sectional. Inferring within-subject decline and aging from cross-sectional associations in people of different-age cohorts has many pitfalls and is prone to confuse aging with cohort differences (e.g., IQ scores have increased in members of more recent cohorts, and there are marked generational differences in exposure to diseases, toxins, antibiotics, education, and nutrition which can influence brain measures, including neuroimaging data)^17–19^. Cross-sectional observations that older brain-age is associated with ADRD and many of its risk factors are consistent with at least two perspectives on brain aging, each of which has distinct implications.

The first perspective is that older brain-age could be an indicator of accelerated brain aging that has accumulated over an individual’s lifetime and increases susceptibility to ADRD and age-related cognitive decline. This perspective implies that at some point in early development, all individuals have a brain-age that is young and the same as their young chronological age. Brain-age scores then diverge with time from chronological age, as genetic, environmental, and lifestyle factors create variation in the rate of brain aging. Here we will refer to this perspective broadly as the “geroscience perspective”^20^. This perspective is based on the geroscience hypothesis which states that aging is the result of deterioration across multiple organ systems and that furthermore this deterioration is the root cause of age-related disease. It is hypothesized that treatments that can slow this decline will therefore reduce the risk for age-related disease. This theoretical interpretation of brain-age is the dominant interpretive framework found in the brain-age literature^10,11,21^.

The second perspective on brain aging is the “early system integrity” perspective of cognitive/biological aging^22^. According to this perspective, individuals vary in their brain and body health from the beginning of life. Moreover, according to the system-integrity view, the correlation between brain and body health persists across the lifespan so that both brain and body health predict aging outcomes^23–25^. From this perspective, the reason brain-age predicts ADRD and mortality later in life is because brain-age is an indicator of compromised lifelong brain health^26,27^. Instead of reflecting accelerated brain aging and the brain’s accumulated biological degeneration, an older brain-age at midlife reflects compromised system integrity that has been present since childhood and stable for decades.

Here we tested whether an older brain-age is associated with accelerated aging or reflects stable individual differences in system integrity in the Dunedin Longitudinal Study. First, we hypothesized that if individuals with an older brain-age have brains that are aging faster, they should also have a body that has aged faster, given that, according to the geroscience perspective, aging is the progressive, generalized deterioration and loss-of-function across multiple organ systems^28,29^. Second, we hypothesized that if individuals with older brain-age have undergone accelerated aging they should show signs of cognitive decline^30^. In contrast, if older midlife brain-age represents system integrity from early life, we hypothesized that older brain-age should be correlated with poorer neurocognitive functioning as assessed already in early childhood.

## Methods

### See Supplement for Expanded Methods

#### Participants

Participants are members of the Dunedin Longitudinal Study, a representative birth cohort (N = 1,037; 91% of eligible births; 52% male) born between April 1972 and March 1973 in Dunedin, New Zealand (NZ), who were eligible based on residence in the province and who participated in the first assessment at age 3 years^31^. The cohort represented the full range of socioeconomic status (SES) in the general population of NZ’s South Island and as adults matches the NZ National Health and Nutrition Survey on key adult health indicators (e.g., body mass index (BMI), smoking, GP visits) and the NZ Census of citizens of the same age on educational attainment. The cohort is primarily white (93%), which matches the demographics of the South Island. Assessments were carried out at birth and ages 3, 5, 7, 9, 11, 13, 15, 18, 21, 26, 32, 38, and most recently (completed April 2019) 45 years, when 94% (N = 938) of the 997 participants still alive took part. Each participant was brought to the research unit for 1.5 days of interviews and examinations. Written informed consent was obtained from participants and study protocols were approved by the NZ Health and Disability Ethics Committee. Brain imaging was carried out at age 45 years for 875 study members (93% of age-45 participants). Data from 6 study members were excluded due to major incidental findings or previous head injuries (e.g., large tumors or extensive damage to the brain). This resulted in brain-imaging data for our current analyses from 869 study members, who represented the original cohort (attrition analysis in supplement).

#### MRI Acquisition

Study participants were scanned using a Siemens Skyra 3T scanner (Siemens Healthcare, Erlangen, Germany) equipped with a 64-channel head/neck coil at the Pacific Radiology imaging center in Dunedin, New Zealand. High resolution structural images were obtained using a T1-weighted MP-RAGE sequence with the following parameters: TR = 2400 ms; TE = 1.98 ms; 208 sagittal slices; flip angle, 9°; FOV, 224 mm; matrix = 256×256; slice thickness = 0.9 mm with no gap (voxel size 0.9×0.875×0.875 mm); and total scan time = 6 min and 52 s.

#### Brain-age

We generated brain-age scores using a recently published, publicly-available algorithm^13^. Unlike some of the other publicly-available brain-age algorithms, this method uses a stacked prediction algorithm based on multiple measures of brain structure including cortical thickness, surface area, and subcortical volume all derived from Freesurfer version 5.3^32^. Test-retest reliability was assessed in 20 Dunedin Study members (mean interval between scans=79 days). The ICC of brain-age was. 81, indicating excellent reliability^33^. Moreover, we chose this algorithm because of its performance in predicting chronological age in independent samples and its sensitivity to age-related cognitive impairment in old age^13^. All regression analyses used brain-age scores (i.e., the difference between an individual’s predicted age from MRI data and their exact chronological age, between birth and the date of the MRI scan).

#### Adulthood Measures of Cognitive Functioning and Accelerated Aging

*Cognitive functioning* at age 45 was assessed with the Wechsler Adult Intelligence Scale-IV^34^, which measures the Intelligence Quotient (IQ) and four specific domains of cognitive function: Verbal Comprehension, Perceptual Reasoning, Working Memory, and Processing Speed. Study members were also tested with an additional suite of measures of vocabulary, memory, and executive functioning (Table 1 and Supplement). *Accelerated aging* was assessed (a) by the Pace of Aging, a longitudinal composite of multiple biomarkers that indexes the integrity of metabolic, cardiovascular, respiratory, kidney, immune, and dental systems, measured at 4 study waves from the cohort members’ 20’s to their mid-40’s, and (b) by independent ratings of Facial Aging. All measures are described in Table 1.

**Tables 1.** Description of Study Measures.

| Measure | Description |
| --- | --- |
| **Adult Cognitive Assessment** |  |
| Adulthood IQ | IQ at age 45 was measured with the Wechsler Adult Intelligence Scale-IV (WAIS-IV)<sup>52</sup>. The WAIS-IV generates the overall full-scale IQ, and four WAIS-IV indexes assess abilities that make up the IQ: Processing Speed, Working Memory, Perceptual Reasoning, and Verbal Comprehension. In addition, we examine performance on the digit symbol substitution<sup>53</sup> subtest, which is most representative of “fluid” cognitive ability, and on the Rey Auditory Verbal Learning test<sup>54</sup> of memory. |
| **Childhood Assessments** |  |
| Childhood Brain Health (age 3) | Age-3 Brain Health is a composite measure from a 45-minute examination that included assessments by a pediatric neurologist, standardized tests of intelligence, receptive language, and motor skills, and examiners’ ratings of each child’s emotional and behavioral regulation<sup>35</sup>. |
| Childhood IQ | IQ was measured with the Wechsler Intelligence Scale for Children-Revised (WISC-R<sup>36</sup>; averaged across ages 7, 9, and 11). In addition, we examine performance on the digit-span subtest and, the Rey Auditory Verbal Learning test<sup>54</sup> of memory. |
| **Measures of Accelerated Aging** |  |
| Pace of Aging | Pace of Aging was measured for each Dunedin Study member with repeated assessments of a panel of 19 biomarkers taken at ages 26, 32, 38, and 45 years (see supplemental for details), as previously described<sup>6</sup>. |
| Facial Aging | Facial Aging was based on ratings by an independent panel of 8 raters of each Study member’s facial photograph. |

**Table 2.** Associations Between Brain-Age at 45 Years, Measures of Cognitive Functioning, Accelerated Aging, and Cognitive Decline.

| **Variable** | **n** | **Standardized $\beta$ (95% CI)** | **P value** |
| --- | --- | --- | --- |
| **Adulthood Cognitive Function, age 45** |  |  |  |
| IQ | 867 | -0.20 (-0.27 to -0.14) | <0.001 |
| Processing speed | 867 | -0.12 (-0.19 to -0.05) | <0.001 |
| Working memory | 864 | -0.15 (-0.22 to -0.09) | <0.001 |
| Perceptual reasoning | 867 | -0.17 (-0.23 to -0.10) | <0.001 |
| Verbal comprehension | 857 | -0.19 (-0.26 to -0.13) | <0.001 |
| RAVL memory test (Total score) | 867 | -0.14 (-0.21 to -0.07) | <0.001 |
| RAVL memory test (Recall score) | 863 | -0.09 (-0.16 to -0.02) | 0.012 |
| Digit Symbol Coding | 867 | -0.15 (-0.22 to -0.08) | <0.001 |
| **Childhood Cognitive Function** |  |  |  |
| IQ | 859 | -0.18 (-0.24 to -0.11) | <0.001 |
| Performance IQ | 847 | -0.14 (-0.21 to -0.08) | <0.001 |
| Verbal IQ | 847 | -0.17 (-0.24 to -0.11) | <0.001 |
| RAVL memory test (Total score)* | 644 | -0.13 (-0.20 to -0.05) | <0.001 |
| RAVL memory test (Recall score)* | 643 | -0.11 (-0.18 to -0.04) | 0.003 |
| Digit Symbol Coding | 847 | -0.09 (-0.15 to -0.02) | 0.014 |
| **Early Childhood Neurocognitive Status** |  |  |  |
| Age-3 Brain Health | 867 | -0.12 (-0.19 to -0.05) | <0.001 |
| **Biological Aging** |  |  |  |
| Accelerated Pace of aging (age 26 to 45) | 868 | 0.22 (0.15 to 0.28) | <0.001 |
| Facial age (age 45) | 868 | 0.15 (0.09 to 0.22) | <0.001 |
| Accelerated facial aging (age 38 to 45) | 864 | 0.07 (0.02 to 0.12) | 0.009 |
| **Cognitive Decline (childhood to age 45)** |  |  |  |
| IQ | 857 | -0.07 (-0.12 to -0.03) | 0.001 |
| RAVL memory test (Total score)* | 644 | -0.12 (-0.19 to -0.05) | 0.001 |
| RAVL memory test (Recall score)* | 643 | -0.08 (-0.15 to -0.01) | 0.028 |
| Digit Symbol Coding | 804 | -0.10 (-0.15 to -0.04) | <0.001 |

RAVL: Rey Auditory Verbal Learning.
\*Only children attending the research unit were administered the RAVL, resulting a smaller sample size with data on this neuropsychological test.

#### Childhood Measures of Brain Health and Cognitive Functioning

At age 3 years, each child participated in a 45-minute examination that included assessment by a pediatric neurologist and standardized tests of intelligence, receptive language, and motor skills. Afterwards the examiners (having no prior knowledge of the child) rated each child’s emotional and behavioral regulation during the protocol. These 5 measures were combined to yield an index of Brain Health (Table 1 and Supplement)^35^. In late childhood (ages 7, 9, and 11 years), Study members were administered the Wechsler Intelligence Scale for Children–Revised (WISC-R) yielding IQ scores^36^. Scores from the three WISC-R administrations were averaged to yield a single, reliable measure of childhood cognitive function. Study members were also tested with an additional suite of measures of vocabulary, memory, and executive functioning (Table 1).

#### Statistical Analysis

We tested associations between brain-age and all target variables using linear regression models in R (version 3.4.0). All models were adjusted for sex. Cognitive decline from childhood to adulthood was measured using a statistical adjustment approach that tested deviation (or change) in a participants’ adult IQ from what would be expected based on their childhood IQ. The premise and analysis plan for this project were pre-registered on https://sites.google.com/site/dunedineriskconceptpapers/documents. Analyses reported here were checked for reproducibility by an independent data-analyst, who recreated the code by working from the manuscript and applied it to a fresh dataset.

## Results

### People of the same chronological age differ in brain-age

As illustrated in Figure 1, despite the narrow range of chronological ages in the Dunedin Study (mean = 45.15, SD = 0.69, range = 43.48 - 46.98), there was substantial variation in brain-age (mean = 40.93, SD = 8.04, range = 23.84 - 71.63). The slight bias towards lower predicted brain-age in this midlife cohort (i.e., we observe younger mean brain-age than mean chronological age) is consistent with findings in this field of research, where brain-age algorithms appear to systematically overestimate mean brain predicted age before age 35 and underestimate mean brain predicted age after age 35^37^.

**Figure 1.**
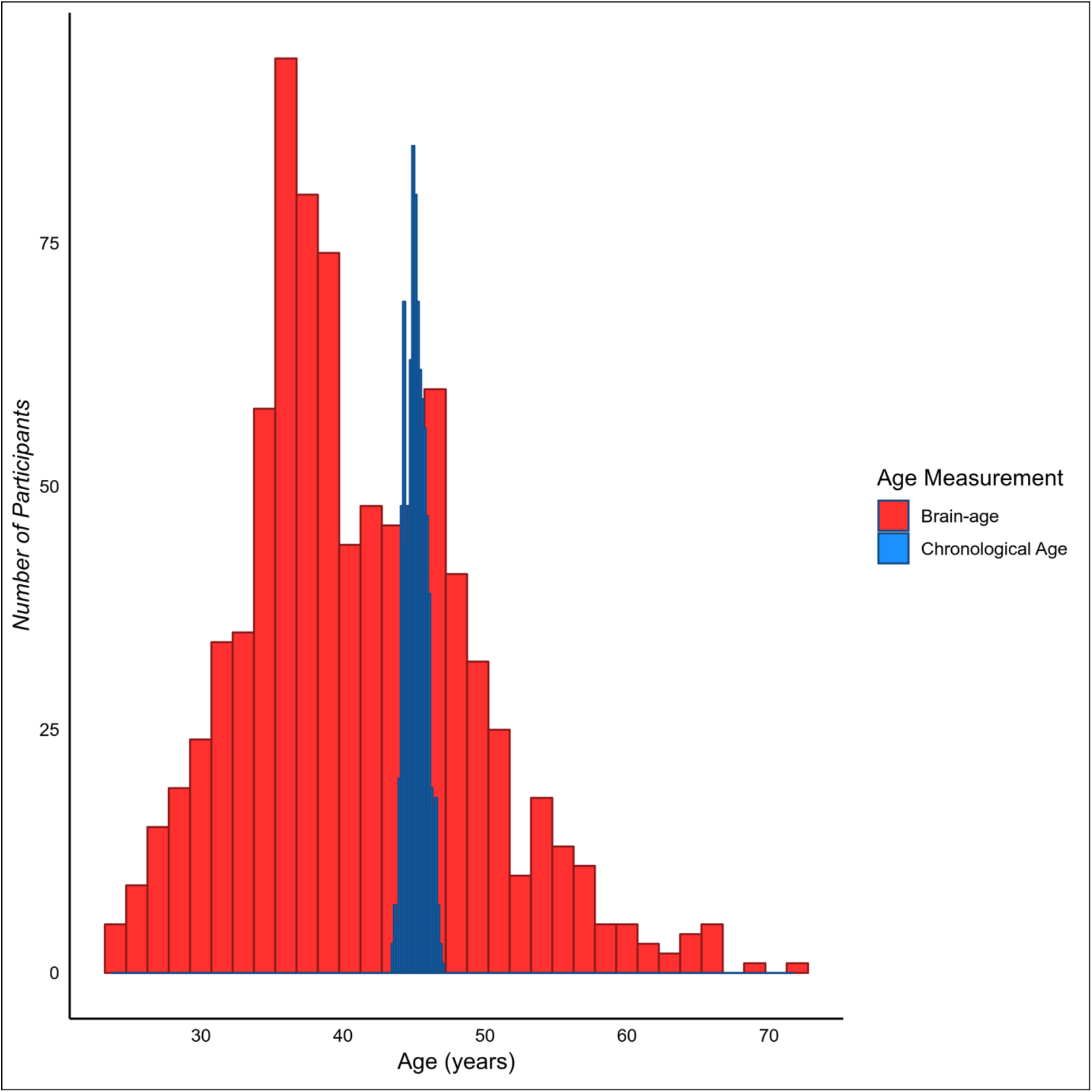
The distribution of chronological age and brain-age amongst the Dunedin Study members. While there is very little variation in chronological age, there is a large amount of variation in brain-age.

### Older brain-age and adult cognitive function

Both the system-integrity and geroscience perspectives predict that brain-age should be associated with cognitive function. Consistent with both perspectives, Study members with older brain-ages performed more poorly on cognitive tests (Table 1). Those with older brain-age had lower full-scale IQ at age 45 (standardized β = −0.20, 95% CI = −0.27 to −0.14; p < .001; Figure 2). However, the associations between brain-age and cognitive functions were non-specific; Study members with older brain-ages had lower scores on all IQ subscales at age 45 including verbal comprehension, which is a crystallized measure (standardized β = −0.19, 95% CI = −0.26 to −0.13; p < .001), and the three fluid measures: perceptual reasoning (standardized β = −0.17, 95% CI = −0.23 to −0.10; p < .001), processing speed (standardized β = −0.12, 95% CI = −0.19 to - 0.05; p < .001), and working memory (standardized β = −0.15, 95% CI = −0.22 to −0.09; p < .001). In addition, Study members with older brain-ages performed more poorly on additional cognitive tests known to be particularly sensitive to age-related deterioration^38^, including digit symbol coding (standardized β = −0.15, 95% CI = −0.22 to −0.08; p < .001), as well as tests of memory (Rey total learning: standardized β = −0.14, 95% CI = −0.21 to −0.07; p < .001; and Rey delayed-recall scores: standardized β = −0.09, 95% CI = −0.16 to −0.02; p = .012).

**Figure 2.**
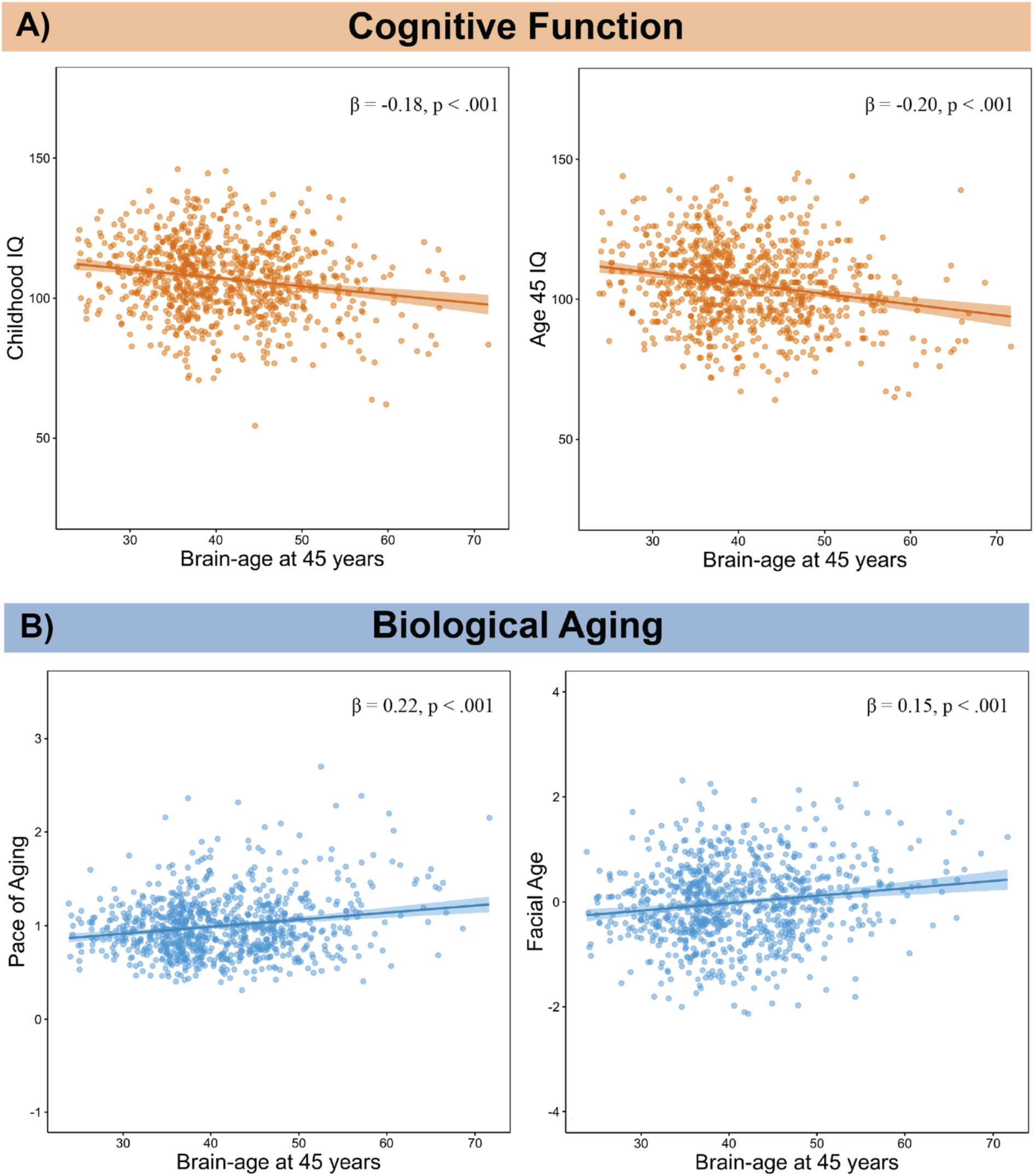
Panel A displays associations between older age-45 brain-age and lower cognitive function. The left panel displays the association between the older brain-age and lower childhood IQ. The right panel displays the association between the older brain-age and lower IQ measured at age 45. Panel B displays associations between older age-45 brain-age and accelerated biological aging. The left panel displays the association between accelerated pace of biological aging between ages 26 and 45 and older brain-age. The Pace of Aging quantifies Study members’ rate of biological aging in year-equivalent units of physiological decline occurring per chronological year. The average Study member experienced 1 year of physiological decline per each chronological year, a Pace of Aging of 1. The right panel displays a scatter plot of the association between older facial age and older brain-age. To illustrate facial aging, the right panel shows digitally averaged faces of the ten male and female Study members rated as looking the oldest and the ten male and female Study members rated as looking the youngest. Facial Age is standardized to M=0, SD=1.

### Older brain-age, childhood cognitive function, and Brain Health

The system-integrity perspective predicts that associations between brain-age and cognitive functions are present since childhood. Consistent with this prediction, 45-year-olds with older brain-age had lower full-scale IQ when measured in late childhood (standardized β = −0.18, 95% CI = −0.24 to −0.11; p < .001; Figure 2). Again we did not find evidence for specificity of this association. Study members with older brain-age had lower performance IQ, a fluid measure (standardized β = −0.14, 95% CI = −0.21 to −0.08; p < .001), and lower verbal IQ, a crystallized measure (standardized β = −0.17, 95% CI = −0.24 to −0.11; p < .001). As in adulthood, study members with older brain-age had poorer performance in childhood on digit symbol coding (standardized β = −0.09, 95% CI = −0.15 to −0.02; p = .014). Those with older brain-age also had poorer performance on measures of memory in childhood (Rey total learning: standardized β = - 0.13, 95% CI = −0.20 to −0.05; p < .001; Rey delayed-recall: standardized β = −0.11, 95% CI = - 0.18 to −0.04; p < .001). Finally, consistent with the system-integrity perspective, Study members with older brain-ages at age 45 had poorer Brain Health measured when they were just 3 years old (standardized β = −0.12, 95% CI = −0.19 to −0.05; p < .001).

### Older brain-age is associated with accelerated biological aging

The geroscience perspective predicts that Study members with older brain-ages should have bodies that are aging at a faster rate. We found evidence to support this account as Study members with older brain-age tended to have a faster Pace of Aging from age 26 to 45 (standardized β = 0.22, 95% CI = 0.15 to 0.28; p < .001; Figure 2). Study members in the oldest decile of brain-age aged 1.17 biological years per chronological year between ages 26 to 45 years, compared to just 0.95 biological years per chronological year for those in the youngest decile. This amounted to 4.22 additional years of biological aging, between ages 26 to 45, for those in the highest brain-age decile. Furthermore, those with older brain-age were rated by independent raters as looking physically older than those with younger brain-age (standardized β = 0.15, 95% CI = 0.09 to 0.22; p < .001; Figure 2). In addition Study members with older brain-age declined faster in their facial age scores between age 38 and 45 (standardized β = 0.07, 95% CI = 0.02 to 0.12; p = .009), suggesting older brain-age predicted a faster pace of facial aging over the course of just 7 years.

### Older brain-age and accelerated cognitive aging

Finally, the geroscience perspective also predicts that Study members with older brain-age should show cognitive decline. Consistent with this perspective, Study members with older brain-age showed initial signs of cognitive decline from their childhood IQ scores to their age-45 IQ scores (standardized β = −0.07, 95% CI = −0.12 to -.03; p = .001; Figure 3). This decline was also found in cognitive tests known to be especially sensitive to aging-related cognitive decline^38^ including digit symbol coding (standardized β = −0.10, 95% CI = −0.15 to −0.04; p < .001) and memory tests (Rey total learning: standardized β = −0.12, 95% CI = −0.19 to −0.05; p < .001; Rey delayed recall: standardized β = −0.08, 95% CI = −0.15 to −0.01; p = .028).

**Figure 3.**
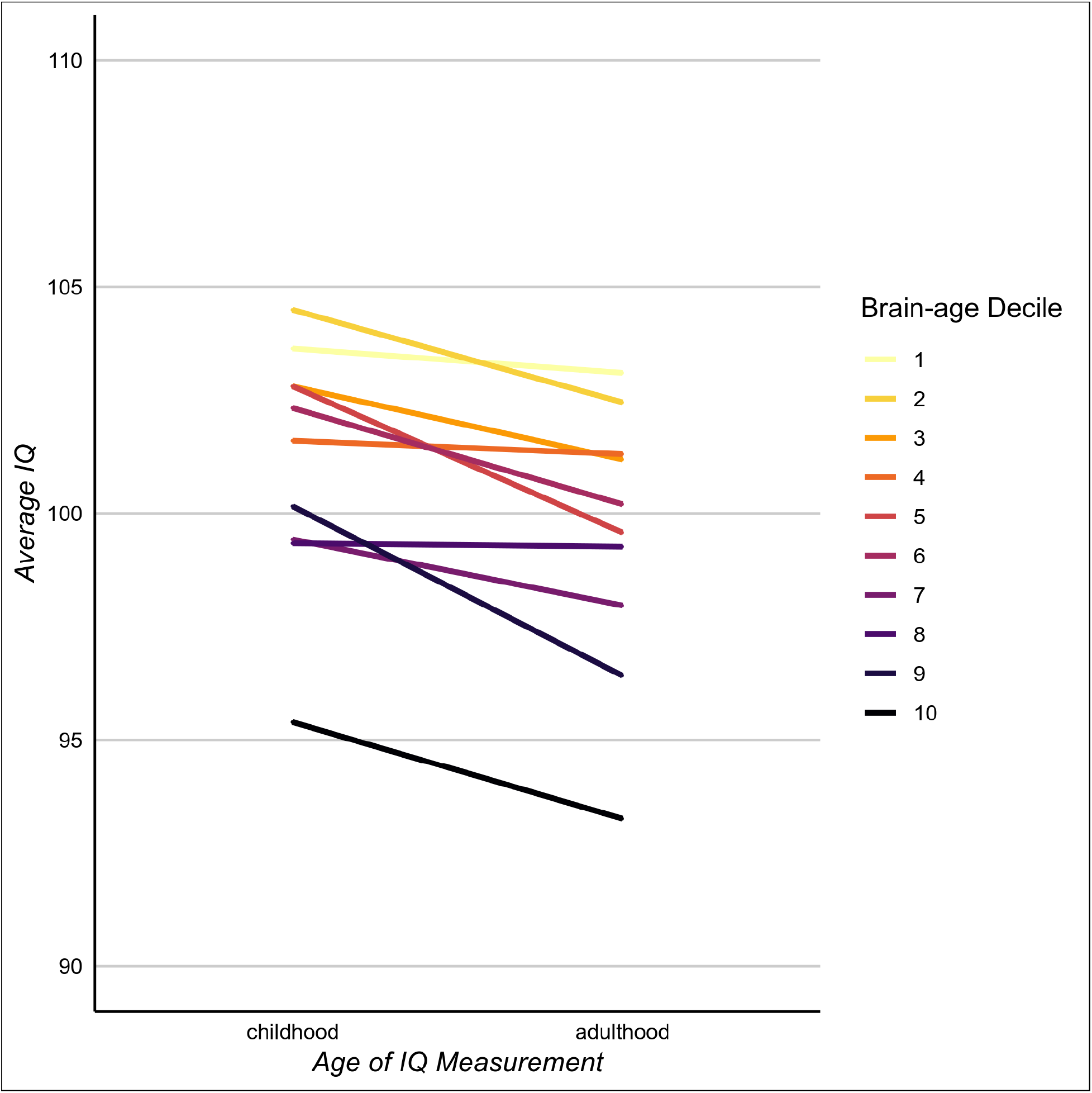
The associations of brain-age with cognitive functioning and cognitive decline. Those with younger age-45 brain-age had the highest IQ scores in both childhood and adulthood. In addition, cognitive decline was greatest among those with older age-45 brain-age; the slopes connecting childhood to adulthood are steeper among Study members with older brain-ages. Sample sizes for each decile from the lowest to the highest WMH volume were: 86, 86, 85, 86, 85, 86, 86, 85, 86, 86.

## Discussion

Using data from a population-representative longitudinal birth cohort followed over four decades, we compared two perspectives of aging (the “geroscience” and “system-integrity” perspectives) that provide disparate explanations for cross-sectional associations between older brain-age and age-related health outcomes (e.g., ADRD and mortality). We found evidence to support both perspectives. Specifically, while Study members with older brain-age had lower cognitive ability in adulthood, they also had poorer cognitive functioning in childhood and poorer brain health already at age three years. These findings are consistent with the system-integrity account of brain-age as representing long-standing brain dysfunction present and stable from early life. However, we also found evidence that individual differences in brain-age were associated with accelerated biological and cognitive aging (e.g., with cognitive decline from childhood to midlife). Together, these findings suggest that an older midlife brain-age is generated by early individual differences (i.e., system-integrity perspective) as well as by accelerated aging that is accumulated throughout a lifetime (i.e., geroscience perspective).

In addition to comparing perspectives of aging, we were able to investigate the relationship between brain-age and aging of the rest of the body. By quantifying each person’s personal pace of biological aging, we were able to demonstrate that Study members with older brain-age had experienced at least two decades of accelerated age-related degradation of the body. Consistent with the “common-cause hypothesis” of aging^28,39,40^, this finding provides evidence that the brain is not exempt from the biological aging that causes a generalized deterioration of organ systems across the body.

A striking finding in research about aging and mortality is that measures of health taken very early in life can predict the likelihood of death and disease much later in life^23^. For example, individuals with low birthweight are at an increased risk for disease and early mortality^29,41^. Consistent with these findings we found that brain-age at age 45 can, in part, be predicted from cognitive function measured in middle childhood and from poor brain health measured at age three years. These findings suggest that accelerated brain deterioration and aging, indexed here with brain-age, may be one mechanism through which individual differences in early system integrity lead to later morbidity and mortality^42,43^. Further research is needed to test whether brain-age mediates the relationship between early deficits in system integrity and later age-related disease.

Our study is not without limitations. First, we do not have childhood brain imaging data that would allow us to directly link accelerated biological aging to accelerated brain aging in the same individuals over time. MRI was not performed in child cohorts during the 1970’s. Previous studies have found that longitudinal changes in brain-age track changes in symptom severity in schizophrenia and cognitive decline in older adults with ADRD^16,44^ but it is not yet known if changes in brain-age track with cognitive decline earlier in the lifecourse.

Second, like other studies of brain-age of which we are aware, the brain-age metric used here was trained on structural MRI data from a large cross-sectional dataset of individuals across a broad age-range^13^. While we have demonstrated that this approach can measure signs of accelerated aging in the brain, it is nevertheless limited in two major ways: 1) brain-age is based on cross-sectional comparisons of individuals of different ages, which do not distinguish cohort effects (cohort differences in exposures) from developmental changes^17,18^. As a result brain-age may be less sensitive to interventions that modify aging processes. 2) Brain-age incorporates only information from T1-weighted structural scans. Diffusion-weighted imaging, fluid-attenuated inversion recovery, and functional imaging are known to change with advancing age and are linked with aging-related brain disease^45–47^. Integrating these additional data types into brain-age algorithms may produce biomarkers more predictive of pathogenic brain aging. Optimal brain-age biomarkers for testing interventions to slow brain aging should be developed from longitudinal, multimodal MRI data that measure accelerated, within-subject brain aging. While many of the effect sizes observed here are modest, such an approach promises to improve the utility of brain-age as a surrogate biomarker for accelerated aging because longitudinal within-subject structural changes have been associated with larger age associations than have cross-sectional comparisons^48^.

Prevention of ADRD is a pressing public health priority due to our rapidly aging population and the lack of effective treatments for ADRD in old age^49,50^. For prevention to be successful, reliable measures are needed of subclinical changes in accelerated brain aging that occur in midlife, decades before the onset of clinically relevant symptoms^3,51^. Such measures would allow accelerated identification of modifiable risk factors, novel treatment targets, and an improved ability to evaluate the effectiveness of preventive interventions. Here we have shown that midlife brain-age is associated with individual differences in the pace of biological and cognitive aging, suggesting that brain-age holds promise as a surrogate biomarker for these purposes, and brain-age measures should continue to be refined. Importantly, we provide evidence that brain-age is a reliable measure in midlife that is indicative of accelerated aging as well as of early system-integrity deficits that may predispose the brain to late-life disease.

## Acknowledgements

This research was supported by National Institute on Aging grants AG032282, AG049789, AG028716, and UK Medical Research Council grant MR/P005918. Additional support was provided by the Jacobs Foundation. MLE is supported by the National Science Foundation Graduate Research Fellowship under Grant No. NSF DGE-1644868. The Dunedin Multidisciplinary Health and Development Research Unit was supported by the New Zealand Health Research Council and MBIE. We thank the members of the Advisory Board for the Dunedin Neuroimaging Study, Dunedin Study members, Unit research staff, and Study founder Phil Silva, PhD, University of Otago.

