## Supplemental Info for "Brain-age in midlife is associated with accelerated biological aging and cognitive decline in a longitudinal birth-cohort"

*Age-3 Brain Health*

At age 3 years, each child in the cohort participated in a 45-minute evaluation that included examination by a pediatric neurologist, plus standardized tests of intelligence (Peabody), receptive language (Reynell), and motor skills (Bayley). Afterwards study staff (having no prior knowledge of the child) rated each child's behavioral self-regulation. Using this information, we created a summary factor score via confirmatory factor analysis which we termed Brain Health, a global index of the child's early neurocognitive status[1,2]. This Brain Health measure was used in our current analyses as a measure of neurological functioning and brain integrity at age 3.

*Pace of Aging (age 45)*

Pace of Aging was measured for each Dunedin participant with repeated assessments of a panel of 19 biomarkers taken at ages 26, 32, 38, and 45 years, as previously described[3]. The 19 biomarkers were: body mass index, waist-hip ratio, glycated hemoglobin (HbA1C), leptin, blood pressure (mean arterial pressure), cardiorespiratory fitness (VO2Max), forced expiratory volume in one second (FEV1), forced vital capacity ratio (FEV1/FVC), total cholesterol, triglycerides, high-density lipoprotein (HDL) cholesterol, apolipoprotein B100/A1 ratio, lipoprotein(a), creatinine clearance, urea nitrogen, C-reactive protein, white blood cell count, gum health, and caries-affected tooth surfaces. Measures were taken in counterbalanced order across participants with the exception of blood, which was drawn at the same time of day for all participants at all four ages and dental examinations which were conducted in the late afternoon at all four ages. Women who were pregnant at the time of a given assessment were excluded from that wave of data collection. The measurement of each biomarker is described below. Change over time in each biomarker was modeled with mixed-effects growth models, and these rates of change were combined into a single index scaled in years of physiological change occurring per one chronological year, as per the method previously described[3]. Participants ranged in their Pace of Aging from near 0 y of physiological change per chronological year to nearly 3 y of physiological change per chronological year.

| | |
|---|---|
| ***Body mass index*** | Height was measured to the nearest millimeter using a portable stadiometer (Harpenden; Holtain, Ltd.). Weight was measured to the nearest 0.1 kg using calibrated scales. Individuals were weighed in light clothing. Body mass index (BMI) was calculated. |
| ***Waist-hip ratio*** | Waist girth was the perimeter at the level of the noticeable waist narrowing located between the costal border and the iliac crest. Hip girth was taken as the perimeter at the level of the greatest protuberance and at about the symphysion pubic level anteriorly. Measurements were repeated and the average used to calculate waist-hip ratio. |
| ***Glycated hemoglobin (HbA1C)*** | Whole blood glycated hemoglobin concentration (expressed as a percentage of total hemoglobin) was measured by ion exchange high performance liquid chromatography (Variant II: BioRad, Hercultes, Calif.), a method certified by the US National Glycohemoglobin Standardization Program (http://www.ngsp.org/). |
| ***Leptin*** | Serum leptin (μg/L) was measured using the Quantikine ELISA Human Leptin Immunoassay (Cat# SLP00, R&D Systems Inc, Minneapolis, MN) according to the manufacturer's instruction. |
| ***Blood pressure (mean arterial pressure)*** | Systolic and diastolic blood pressure were assessed according to standard protocols with a Hawksley random-zero sphygmomanometer with a constant deflation valve. Mean arterial pressure (MAP) was calculated using the formula Diastolic Pressure+1/3(Systolic Pressure - Diastolic Pressure). |

| | |
|---|---|
| ***Cardiorespiratory fitness ($VO_2Max$)*** | Cardiorespiratory fitness was assessed by measuring heart rate in response to a submaximal exercise test on a friction-braked cycle ergometer. Dependent on the extent to which heart rate increased during a 2-min 50 W warm-up, the workload was adjusted to elicit a steady heart-rate in the range 130–170 beats per minute. After a further 6-min constant power output stage, the maximum heart rate was recorded and used to calculate predicted maximum oxygen uptake adjusted for body weight in milliliters per minute per kilogram ($VO_2$max) according to standard protocols[4]. |
| ***Lung function ($FEV_1$ and $FEV_1/FVC$)*** | We calculated post-albuterol forced expiratory volume in one second ($FEV_1$) and the ratio of $FEV_1$ to forced vital capacity (FVC; $FEV_1/FVC$) using measurements from spirometry conducted with a Sensormedics body plethysmograph (Sensormedics Corporation, Yorba Linda, CA, USA). |
| ***Total cholesterol, triglycerides, and high-density lipoprotein (HDL) cholesterol*** | Serum non-fasting total cholesterol, triglycerides, and high-density lipoprotein (HDL) cholesterol levels (mmol/L) were measured by colorimetric assay on a Hitachi 917 analyzer (ages 26-32), a Modular P analyzer (age 38), and a Cobas c702 analyzer (age 45). |
| ***Apolipoprotein B100/A1 ratio*** | Serum apolipoprotein A1 and apolipoprotein B100 (g/L) were measured by immunoturbidimetric assay on a Hitachi 917 analyzer (ages 26-32), a Modular P analyzer (age 38), and a Cobas c502 (age 45), and the ratio between the two was calculated. |
| ***Creatinine clearance*** | Serum creatinine (mmol/L) was measured by kinetic colorimetric assay on a Hitachi 917 analyzer (age 32), Modular P analyzer (age 38), and Cobas c702 (age 45) (Roche Diagnostics, Mannheim, Germany). For Pace of Aging analysis, creatinine was measured as creatinine clearance, calculated using the National Kidney Foundation CKD-EPI Creatinine Equation (2009)[5]. |
| ***Urea nitrogen*** | Serum urea nitrogen (mmol/L) was measured by kinetic UV assay at ages 26 (Hitachi 917 analyzer) and 45 (Cobas c702 analyzer), and by kinetic colorimetric assay at ages 32 (Hitachi 917 analyzer) and 38 (Modular P analyzer). |
| ***High sensitivity C-reactive protein (hsCRP)*** | Serum C-reactive protein (mg/L) was measured by high sensitivity immunoturbidimetric assay on a Hitachi 917 analyzer (age 32), a Modular P analyzer (age 38), and a Cobas c702 (age 45). Values were log-transformed for analysis. |

| | |
|---|---|
| ***White blood cell count*** | Whole blood white blood cell counts ($x10^9$/L) were measured by flow cytometry with a Coulter STKS (Coulter Corporation, Miami, FL) (age 26), a Sysmex XE2100 (Sysmex Corporation, Japan) (age 32), and a Sysmex XE5000 (Sysmex Corporation, Japan) (ages 38 and 45). Counts were log-transformed for analysis. |
| ***Gum health (combined attachment loss)*** | Calibrated dentists examined periodontal health at three sites (mesiobuccal, buccal, and distolingual) per tooth[6]. Gingival recession (the distance in millimeters from the cementoenamel junction to the gingival margin) and probing depth (the distance from the probe tip to the gingival margin) were recorded using a PCP-2 periodontal probe (Hu-Friedy; Chicago). The combined attachment loss for each site was computed by summing gingival recession and probing depth (third molars were not included) and then averaged across all periodontally examined teeth. |
| ***Caries-affected tooth surfaces*** | Teeth were examined for caries and restorations following the World Health Organization Oral Health Surveys methodology. Four surfaces were considered for anterior teeth (canines and incisors): buccal, lingual, distal, and mesial; a fifth surface, occlusal, was considered for premolar and molar teeth. Tooth surfaces were classified as having untreated caries (DS) if a cavitated carious lesion was present, as filled (FS) if a dental restoration was present (including crowns), and missing due to caries (MS) if the participant indicated that a given tooth had been removed due to decay or failed dental restorative work. DS, MS, and FS counts were summed (ranging from 0 to 148 surfaces). Surfaces of teeth that were unerupted, lost due to trauma, extracted for reasons other than caries (impaction, orthodontic treatment, or periodontal disease), or could not be visualized by the examiner were excluded from calculations. |

### *Facial Aging (age 45)*

Facial Aging was based on ratings by an independent panel of 8 raters of each participant's facial photograph. Facial Aging was based on two measurements of perceived age. First, Age Range was assessed by an independent panel of 4 raters, who were presented with standardized (non-smiling) facial photographs of participants and were kept blind to their actual age. Raters used a Likert scale to categorize each participant into a 5-year age range (i.e., from 20-24 years old up to 70+ years old) (interrater reliability = .77). Scores for each participant were averaged across all raters. Second, Relative Age was assessed by a different panel of 4 raters, who were told that all photos were of people aged 45 years old. Raters then used a 7-item Likert scale to assign a "relative age" to each participant (1=”young looking”, 7=”old looking”) (interrater reliability = .79). The measure of perceived age at 45 years, Facial Age, was derived by standardizing and averaging Age Range and Relative Age scores.

**Phase 45 Attrition Analysis**

We conducted an attrition analysis using childhood intelligence quotient (IQ; Supplementary Fig.1) and socioeconomic status (SES; Supplementary Fig. 2) to determine whether participants in the Phase 45 data collection were representative of the original cohort.

Attrition Analysis of Childhood IQ in Phase 45

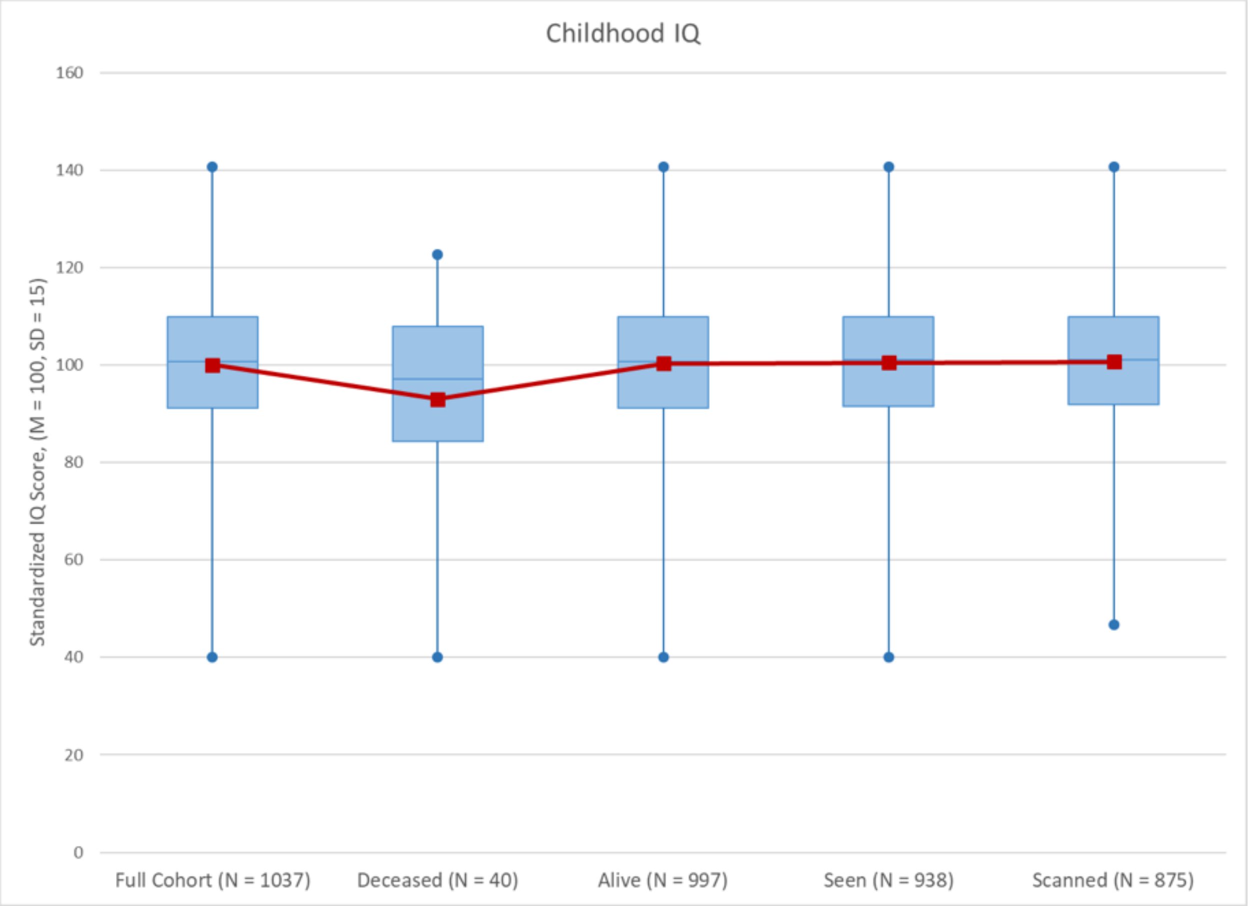


**Supplementary Figure 1**. No significant differences in childhood IQ were found between the full cohort, those still alive, those seen at Phase 45 or those scanned at Phase 45. Those who were deceased by the Phase 45 data collection had significantly lower childhood IQ's than those who were still alive ($t = 2.09$, $p = 0.04$).

Attrition Analysis of Childhood SES in Phase 45

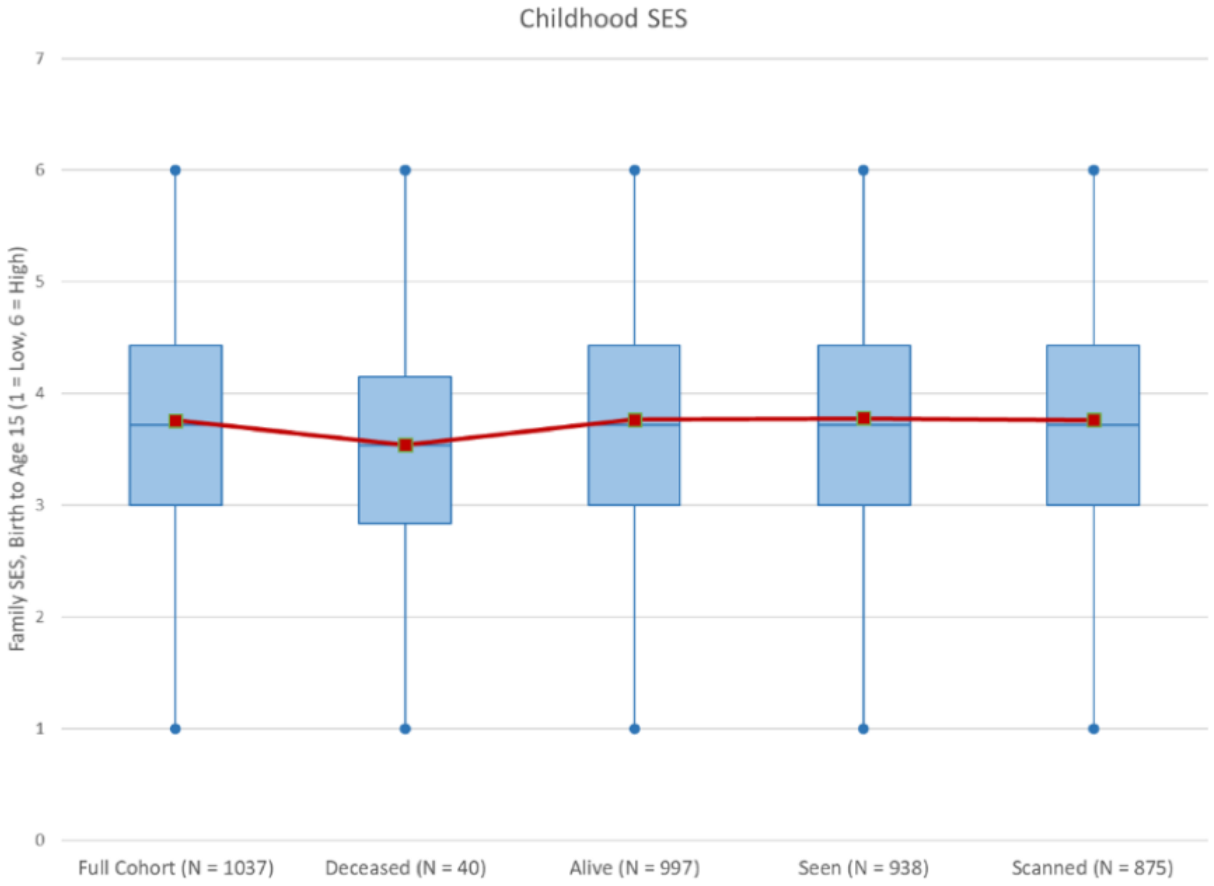


**Supplementary Figure 2.** No significant differences were found between the full cohort, those deceased, those alive, those seen at Phase 45 or those scanned at Phase 45 on childhood SES.

6 World Health Organization. *Oral Health Surveys: Basic Methods*. World Health Organization, 2013.
